# Architecture, Activation, and Conformational Plasticity in the GluA4 AMPA Receptor

**DOI:** 10.1101/2025.06.12.659357

**Authors:** W. Dylan Hale, Richard L. Huganir, Edward C. Twomey

## Abstract

AMPA-subtype glutamate receptors (AMPARs), composed of subunits GluA1-4, mediate fast, excitatory synaptic transmission in the brain. After glutamate binding, AMPAR ion channels exhibit multiple subconductance states that tune neuronal responses to glutamate. GluA4 is the rarest subunit in the brain but is enriched in interneurons. Rising evidence points to the role of GluA4 AMPARs in the development of neurological diseases, but the structural mechanisms of GluA4 function have remained enigmatic. Here, from bilayer recordings and cryo-electron microscopy (cryo-EM), we report the unique features of GluA4 AMPARs that tune receptor function. We find that GluA4 AMPARs have a canonical “Y” shaped architecture where local dimer pairs are domain-swapped between the amino terminal domain (ATD) and ligand binding domain (LBD), both of which comprise the extracellular domain. All four LBDs are glutamate bound yet open the GluA4 ion channel by asymmetric hinging in all channel helices. We observe that the glutamate-saturated LBD has conformational plasticity, and the different conformations of the LBD tune the ion channel gate below. These data provide a framework for understanding how channel subconductance can occur during conditions of saturating glutamate, outline the unique properties of GluA4, expand our understanding of conformational plasticity in AMPARs, and will inform therapeutic design.

---

Fast, excitatory neurotransmission in the vertebrate brain is principally mediated through AMPA-subtype glutamate receptors (AMPARs) – ligand-gated ion channels that open a cation-selective pore in the neuronal membrane in response to binding the neurotransmitter glutamate^1,2^. Transmission through AMPARs is thought to mediate essential processes in cognition, and plasticity of AMPARs alters synaptic strength to mediate the most well-characterized physiological correlates of neurological memory. Dysfunction in AMPAR signaling is increasingly implicated in diverse neurological and psychiatric diseases, including Alzheimer’s dementia, intellectual disability, and schizophrenia^1^. Accordingly, there is intense interest in understanding the mechanisms by which AMPARs function and are regulated at synaptic sites.

AMPARs are tetrameric assemblies formed by four pore-forming subunits from the GluA family of proteins (GluA1-4). AMPAR subunits are unevenly expressed among neuronal cell types and across brain regions^3^, and the subunit composition of AMPARs at synapses can be dynamically regulated during certain forms of AMPAR plasticity^4^. AMPARs bearing the widely expressed GluA2 subunit are rendered impermeable to calcium and structures of native AMPARs purified from brain suggest that GluA2 maintains a privileged structural relationship within the receptor^5^. However, increasing evidence suggests a critical role for calcium-permeable AMPARs (CP-AMPARs) in the brain^6,7^, which either lack GluA2 or include rare calcium-permeable forms of GluA2^8^.

Certain cell types in the brain, such as disease-associated Parvalbumin-expressing interneurons (PVINs) express relatively little of the GluA2 subunit and instead express relatively high levels of the poorly understood calcium-permeable GluA4 subunit^9,10^, which exhibits distinct physiological properties^11–13^. Accordingly, the calcium-permeability of these receptors is critical to the physiological properties of PVINs and altering the calcium permeability in these cells undermines their organismal function^9^. Plasticity of AMPARs on PVINs in cortical cultures can be mediated by GluA4, highlighting the importance of this subunit to the physiology of these cells^14,15^. Rare variants in the gene encoding GluA4 are increasingly found in patients with severe intellectual disability and epilepsy, highlighting the role of GluA4 in human disease^16^. Other than in sparse interneuron classes, GluA4 expression is high in areas of the auditory system^17–22^ and cerebellum^3,11,23,24^, where it likely plays specialized roles in cellular and organismal physiology^25–27^. Dysfunctional GluA4 expression and signaling have also been associated with glioblastoma growth and tumor proliferation^28–30^. However, what mediates the distinct features of GluA4 function is enigmatic.

To address this, we used cryo-EM to solve the structure of a GluA4 homotetrameric AMPAR in complex with the auxiliary subunit TARPγ2 in the presence of glutamate and positive allosteric modulator cyclothiazide (CTZ). We identified a high degree of conformational heterogeneity in the extracellular N-terminal domain (NTD) and ligand binding domain (LBD) of the receptor that may underlie some of the unique functional properties of the receptor and its relevance to cell types such as PVINs. These findings shed light on how GluA4-containing AMPARs differ from other calcium-permeable AMPAR subtypes and the heterogeneity in the LBD layer may provide an alternative model to the current conception of how different subconductance states of AMPARs are achieved.

## Results

### Purification of Functional GluA4 Homotetramers

In the cortex, GluA4 is expressed in PVINs, which also express the auxiliary subunit TARPγ2^31,32^, which has been observed to modulate GluA4 function^33^. Therefore, to solve the structure of a GluA4 homotetrameric AMPAR we employed a fusion construct between GluA4 and TARPγ2 by truncating the GluA4 sequence after the fourth transmembrane domain, followed by a short, flexible linker region, and then beginning the coding sequence for TARPγ2. TARPγ2 was truncated after the TM4 helix, and a thrombin cleavage site, GFP and polyhistidine affinity tag were added to the C-terminus immediately preceding the translational stop. This approach has been used extensively to study AMPAR/TARP complexes^34–38^ and a similar construct has been extensively characterized in whole cell and single-channel preparations^33^. Expression of this construct resulted in a large complex assessed by Fluorescence Size Exclusion Chromatography corresponding to an AMPAR tetramer (**Extended Data 1a**). The GluA4 peak was responsive to variations in lipid composition of the solubilization buffer, with a porcine brain lipid extract demonstrating marked improvements in preserving the tetrameric assembly over DDM/CHS buffer (**Extended Data 1b**). We therefore purified GluA4/TARPγ2 complexes by extracting the protein in a buffer containing DDM and porcine brain lipids and then exchanging the complex into a buffer containing glyco-diosgenin (GDN; **Extended Data 1c**).

To test that GluA4 homotetramers were functional, we performed lipid bilayer recordings of single GluA4 channels passed into reconstituted lipid bilayers in the presence of 1 mM glutamate (Glu) and 100 μM cyclothiazide (CTZ). As compared to bilayers without protein, membranes with GluA4 homotetrameric receptors could be readily observed producing short fluxes in the presence of Glu and CTZ (**Fig. 1a**), consistent with currents that have been previously observed for single AMPAR channels^33,39^,. We therefore concluded that purified GluA4/TARPγ2 complexes were capable of glutamate activation and proceeded with structural analysis of complexes purified under these conditions.

**Figure 1.**
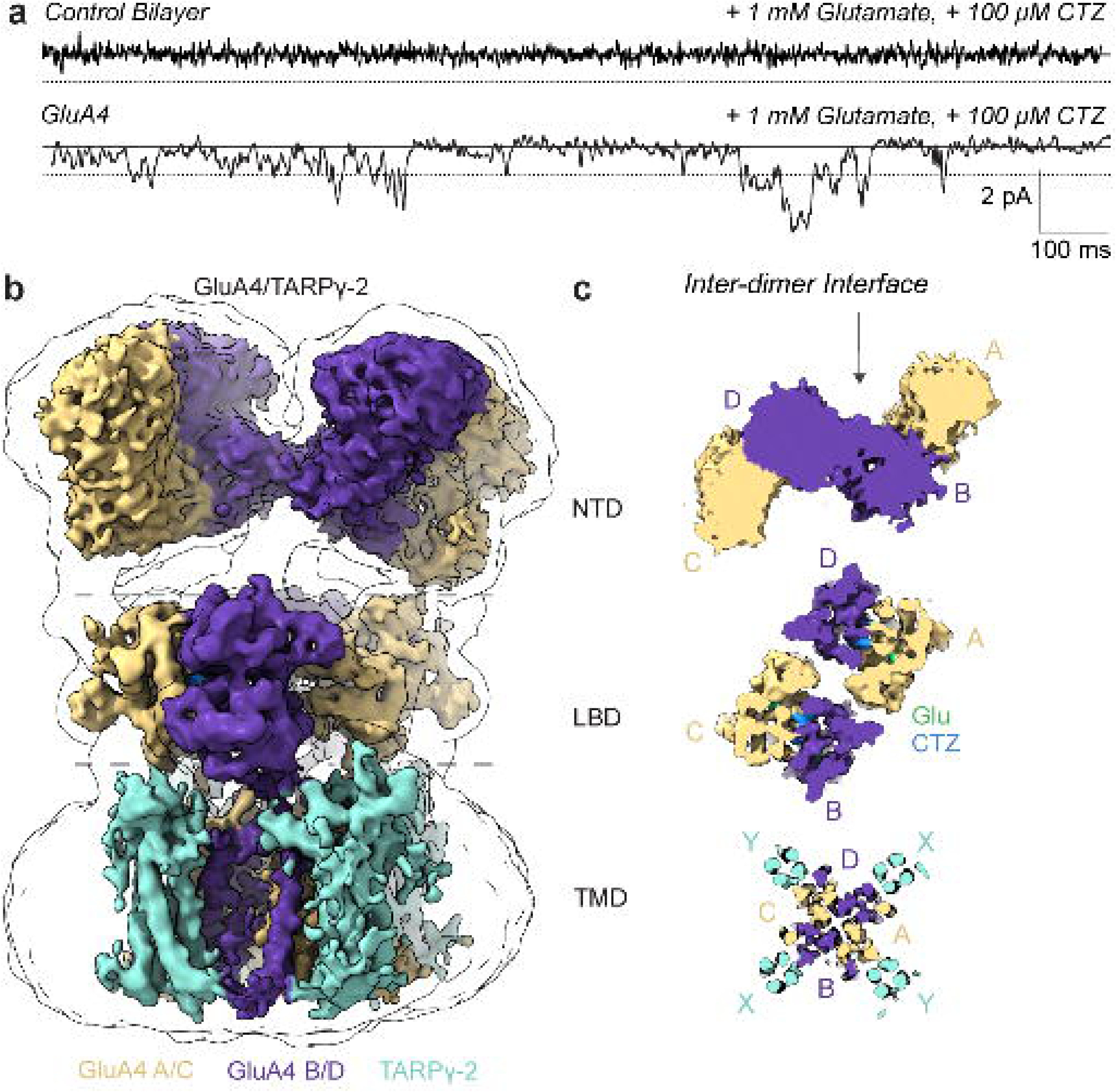
Topology of functional GluA4/TARPγ2 complexes. **a** Single-channel recordings of purified GluA4/TARPγ2 complexes reconstituted in lipid bilayers. The top trace shows the typical responses of bilayers without receptors incubated with 1mM Glu and 100 μΜ CTZ. The bottom trace was recorded with a single GluA4/ΤARPγ-2 complex inserted in the bilayer. **b** Overall topology of GluA4/TARPγ2 complexes showing the preserved Y-shaped AMPAR architecture and an intact NTD. The composite map of the GluA4/TARPγ2 is in color (Extended Data 3, class 8), while a low contour of the whole map is shown as a black outline. **c** Slices through the GluA4/TARPγ2 class 8 composite map showing the intact NTD architecture and domain swapping between the NTD and LBD layers. Density corresponding to Glu and CTZ can be seen within the LBD slices.

### Architecture of GluA4

To examine the structure of activated GluA4, purified GluA4/TARPγ2 sample was spiked with 1 mM Glu and 100 μM CTZ before freezing, and the resulting structure was resolved via CryoEM (**Extended Data 2**). The resulting map displayed a well-resolved LBD and TMD layer, with four TARPγ2s bound around the GluA4 TMD. However, examination of the GluA4/TARPγ2 map revealed that the N-terminal Domain (NTD) was absent from our reconstruction (**Extended Data 2**), suggesting that the NTD of GluA4 may exhibit high conformational heterogeneity, or may even be ruptured as has been recently reported for GluA1/TARPγ3 complexes^40^.

The structural integrity and overall topology of AMPAR complexes depends on AMPAR subunit composition, with GluA2 driving heteromeric assembly of local dimers and contributing to the stabilization of the NTD trans-dimer interface^5,41^. Recent work solving the structure of GluA2-lacking AMPARs has revealed the absence of the trans-dimer interface and a ruptured NTD layer. We therefore wondered whether the NTD was intact in our GluA4/TARPγ2 complex and endeavored to refine the NTD layer of the GluA4 particle set to see if conformational heterogeneity of the NTD layer explained its absence from our reconstruction. 3D classification of the NTD layer alone revealed multiple classes in which NTD lobes could be identified at low resolution (**Extended Data 3a**) and rigid-body fitting of previously determined crystal structures of the GluA4 NTD (PDB: 4GPA, **Extended Data 3b**)^42^, allowed for geometric analysis of the NTD layer.

Density ascribable to four NTDs was observed, with the A subunit NTD forming a local dimer with the B subunit NTD and the C subunit NTD forming a local dimer with the D subunit NTD, as has been previously described for AMPARs. Critically, while significant conformational heterogeneity exists within the GluA4 NTD, the trans-dimer interface between the B and D subunits appears to be preserved (**Fig. 1b,c, Extended Data 3b**), and all resolvable classes of the GluA4 NTD appear to swap local dimer partners below the NTD layer, with the A subunit switching from a local dimer with the B subunit at the level of the NTD to forming a local dimer with the D subunit at the level of the LBD, and the C subunit switching from a local dimer with the D subunit at the level of the NTD to a local dimer with the B subunit at the layer of the LBD (**Fig. 1b**). This contrasts with the NTD arrangement in GluA1 homotetramers where the NTD layer is ruptured with occasional non-domain-swapped GluA1 homotetramers^40^.

To measure the degree of conformational heterogeneity within the NTD layer, we refined maps of the LBD and TMD corresponding to the particle sets contributing to each NTD class (**Extended Data 3a**). Alignment of these maps resulted in composite maps in which differences between the classes in NTD conformation relative to the LBD and TMD layers could be observed. The NTD of GluA4 maintained an intact dimer-of-dimer architecture with the two local NTD dimer pairs angled between 42° and 46° relative to one another (class 9 and class 6, respectively; **Extended Data** Fig. 3c). The NTD remains intact in all resolvable classes but exhibits up to 14° of tilt relative to the pore axis. Furthermore, the distance between the bottom of the NTD and the top of the LBD layer was variable among the classes, exhibiting a 15 Å difference perpendicular along the pore axis (**Extended Data 3c).** Despite these differences in the NTD layer, differences in the overall conformation of the LBD and TMD layers were not observable. Taken together, these findings indicate that the NTD layer of GluA4 homotetramers is intact but exhibits an extraordinary degree of conformational heterogeneity.

### Activation of GluA4

The degree of conformational heterogeneity that we observed in the NTD of activated GluA4 homotetramers suggests that the NTD may be able to act independently of the LBD and TMD layers to carry out functions distinct from channel gating. We therefore examined the activation mechanism of our purified GluA4 homomers, which were prepared in the presence of 1 mM Glu and 100 μM CTZ. Further processing of our consensus particle set produced a map of GluA4 homotetramers containing the LBD and TMD layers at an overall resolution of 3.31 Å, with local resolution below 3Å in the channel pore (**Extended Data 4**). From this map, we observed all four LBD clamshell domains in the closed conformation with Glu bound in the LBD clamshell and CTZ present between the LBD domains (**Fig. 2a, Extended Data 5**). Lipid densities could be resolved around the TMD of the GluA4 assembly that appeared distinct from the previously reported cholesterol densities associated with AMPAR/TARP complexes (**Extended Data 6**), although resolution limitations preclude assignment of lipid identities with high confidence.

**Figure 2.**
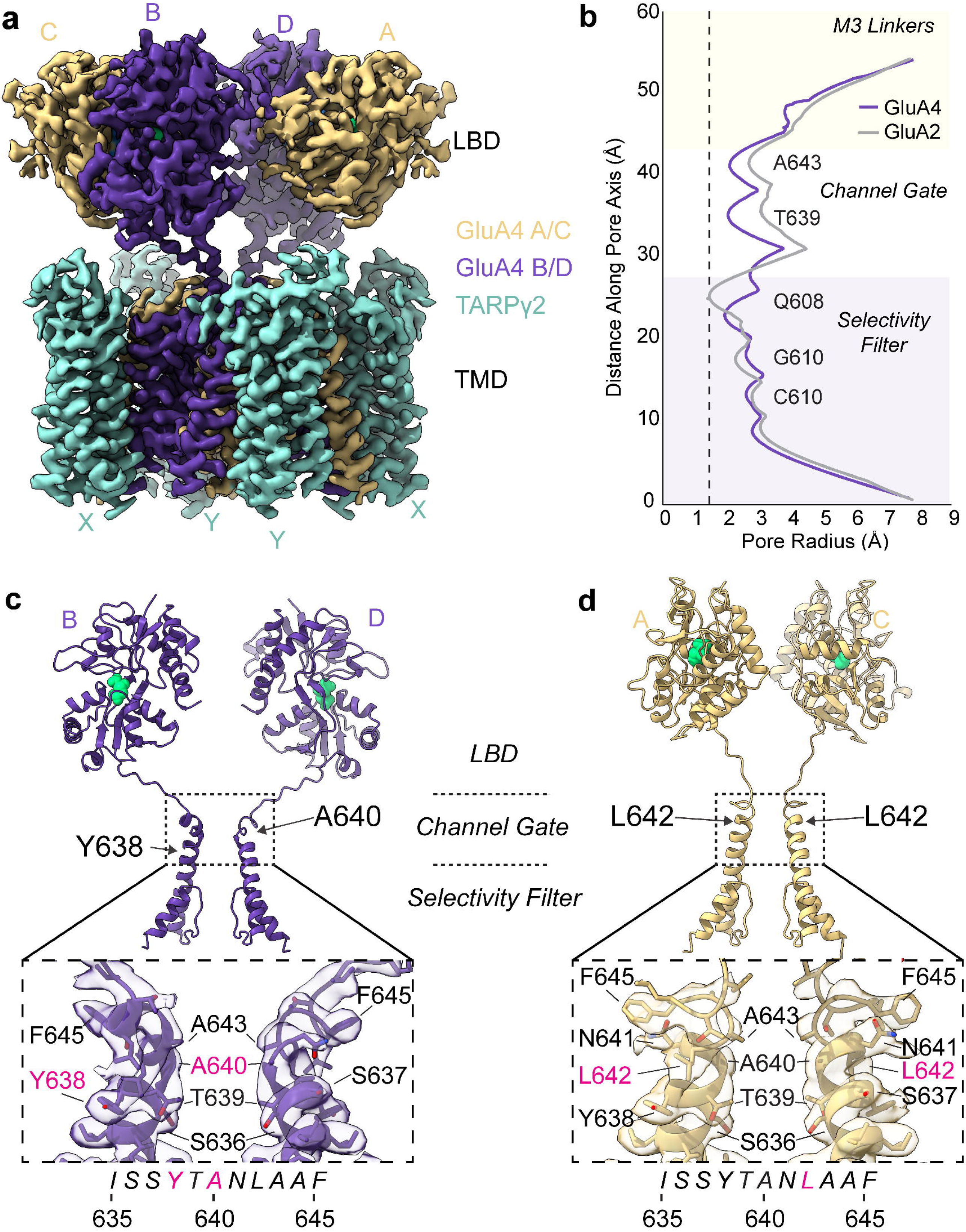
Activation and gating mechanism in GluA4/TARPγ2. **a** consensus map of the GluA4/TARPγ2 LBD and TMD at an overall resolution of 3.31 Å. B and D subunit positions are in purple and A/C subunit positions are in tan. TARPγ2 is shown in teal. Glu bound in the LBD can bee seen in green. **b** Pore radius map of the GluA4/TARPγ2 pore computed using HOLE. the GluA4 pore radius is shown in purple. The radius of a previously solved structure of activated GluA2 is shown in gray (PDB: 5weo). The dashed line indicates the radius of a water molecule (1.4 Å). **c** Activation of GluA4 at the B/D subunit positions. Cutaway of the GluA4/TARPγ2 model at the B and D positions (purple) showing glutamate bound in the LBD (green spheres) and kinking of the M3 helices in the transmembrane domain. Inset shows the map fit at the critical hinging region around A640. **d** The A/C subunit positions in GluA4 (tan) do not show appreciable kinking despite also being bound to Glu in the LBD (green spheres). Inset shows map fit around the upper channel gate.

As in other AMPAR structures, gating in the GluA4 active state is defined by kinking of the M3 helix in the B and D subunits within the “ISSYTANLAAF” motif that couples the top of the M3 to the lower D2 lobe of the ligand binding domain. To confirm that the pore is open in our GluA4 active state, we measured the radius of the ion channel along its z axis. The GluA4 active state measured 2.0 Å at the most constricted point of the upper channel gate (T639 in GluA4, T638 in WT GluA2), indicating an open channel (**Fig. 2b**). The location of this constriction differs from the position of the GluA2 upper channel gate by a single helical turn, (A643 in GluA4, A621 in GluA2). Consistent with recent work showing hinging of the A and C subunit M3 helices during AMPAR activation, the GluA4 active state exhibits asymmetric kinking at the upper channel gate with kinks at all four helices^43^. The B subunit kinks at Y638, and the D subunit kinks at A640 (**Fig. 2c**), while the A and C subunits both kink at L642 (**Fig. 2d**). Together, these data indicate that the conformational heterogeneity in the GluA4 NTD does not meaningfully limit the ability of the GluA4 LBD to activate the channel.

### Conformational Plasticity in the GluA4 LBD

Our structure of the GluA4/TARPγ2 complex revealed the activated state. However, close examination of single channel currents from purified GluA4/TARPγ2 complexes exposed to the same saturating concentrations of Glu and CTZ indicated the presence of multiple distinct levels of conductance through GluA4/TARPγ2 channels (**Fig. 1a, 3a**). Analysis of recordings from several individual channels revealed a distribution of conductances consistent with previous reports of four discreet conductance states for single GluA4/TARPγ2 channels (**Fig. 3a**)^33^. A frequency histogram charting the amplitude of channel openings could be best fit with three gaussian curves, each corresponding to a different magnitude of channel opening. O1 was the dominant open state (8 pS) followed by O2 (18 pS) and then O3 (31 pS). Openings to O4 (45 pS) were observed in the raw traces but were not frequent enough to alter the shape of the frequency histogram. These conductances compare favorably to conductance states recorded from a similar GluA4/TARPγ2 fusion construct using single channel patch-clamp electrophysiology^33^. Given that these subconductance states were sampled by purified receptors under the same conditions that produce our active state structure, where all LBDs are glutamate bound, we wondered how subconductance states might be achieved within a single population of active state receptors.

**Figure 3.**
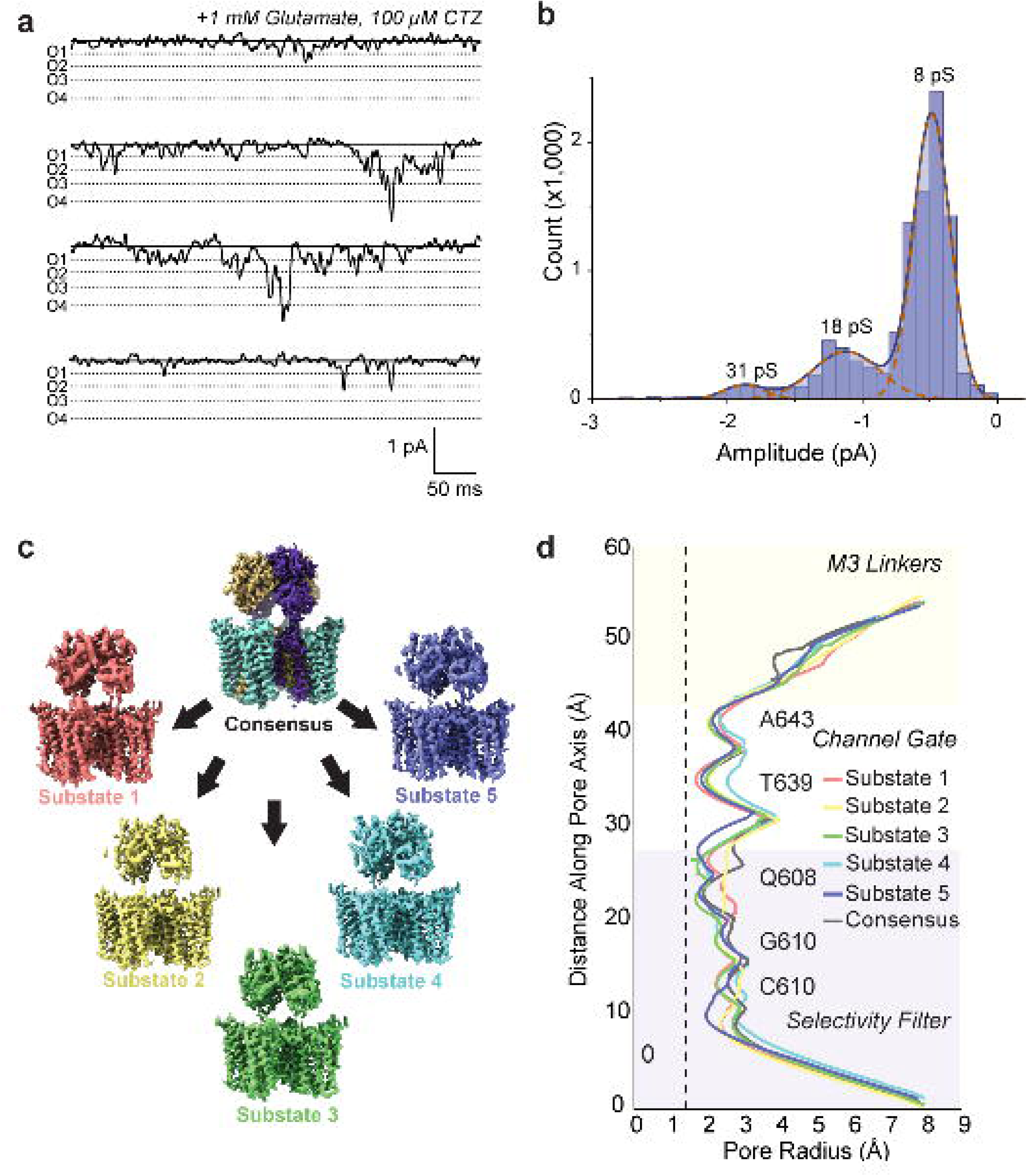
Subconductance and Conformational Heterogeneity in GluA4/TARPγ2. **a** Representative 500ms single channel traces of GluA4/TARPγ2 containing at least one opening event. Dashed lines represent observed conductance levels. **b** Frequency histogram of. Channel openings of GluA4/TARPγ2 showing the fit of the histogram by three gaussian curves with means corresponding to 8 pS, 18 pS and 31 pS, respectively. n = 10 individual channels. **c** 3D Variability analysis was used to generate five independent maps from the consensus particle set composed of non-overlapping particle sets. Each map represents a distinct substate within the consensus particle set. **d** Pore radius plot for each of the five substates, showing variability in the upper channel gate and elsewhere along the channel pore. **e** Overlaid models of substate 1 (red) and substate 5 (blue) demonstrating the degree of change in these substates. The entire LBD layer rotates 7° about the pore axis between substate 1 and substate 5. The entire LBDs also rotate downwards about 18° in the transition between substate 1 and substate 5. These conformations occur despite all LBDs maintaining a closed conformation.

Analysis of the local resolution map of GluA4/TARPγ2 revealed that the local resolution in the LBD layer was noticeably lower than that of the pore-forming TMD, which can sometimes indicate conformational heterogeneity within the dataset (**Extended Data 4**). We therefore used 3D Variability Analysis (3DVA) in CryoSPARC to identify components of variability within our consensus particle set (**Extended Data 7**) that may explain how variable conductances might arise within a population of active state receptors. 3DVA revealed one prominent variability component which appeared to show conformational heterogeneity manifesting as a ‘rocking’ or ‘twisting’ of the LBD layer relative to the TMD layer (**Extended Movie 1**). We therefore separated the particles in our consensus dataset based on where they fell along this variability component (Variability Component 1). We were able to bin the particles into 5 individual groups and to generate 5 independent maps representing different ‘substates’ of the active state receptor (**Fig. 3c**) These substate maps varied in the number of contributing particles and their overall resolution but were each sufficiently well resolved to build five individual models each representing a different LBD/TMD relationship (**Extended Data 8, Extended Data 9**). Since opening of the AMPAR ion channel requires LBD closure to pull open the channel gate, we speculated that variable LBD positions might alter the efficiency of this process and therefore the magnitude of pore opening. We therefore measured the pore radius of each of our solved substates using HOLE^44^ (**Fig. 3d**). We found three sites of variable channel opening between the substates. The most pronounced site was at T639 of the upper channel gate which showed 4 different substates with a lower limit of 1.7 Å and an upper limit of 2.5 Å (**Fig. 3d**).

### GluA4 Substates Exhibit Variable Channel Openings

How are these different substates achieved? Alignment of the TMD layer of the substates revealed overall similar TMD conformations (RMSD = 1.2-1.6 Å) while LBD layer exhibited more heterogeneity (RMSD = 1.3-3.2 Å) **Extended Data 10a**. Notably, each of the substates showed all four LBD clamshells in the closed conformation around a glutamate molecule and CTZ was recognizably present between each LBD local dimer pair (**Extended Data 10b-e**), indicating that the substates did not vary in their ability to bind to the agonist Glu or the positive allosteric modulator CTZ. Close inspection of the substate maps revealed different conformations of the M3 helix critical for channel gating (**Extended Data** Figure 9). Interpolation between the substates revealed a vector of heterogeneity that exhibited variable rotation of the LBD layer with respect to the TMD (**Extended Movie 2)**. This rotation about the pore axis proceeded clockwise from Substate 1 to Substate 5, with 7° of overall rotation of the LBD local dimers (**Fig. 4a**). In addition, all four LBDs rotated downwards away from the pore axis an additional 18° in Substate 5 compared to Substate 1 (**Fig. 4b**). The entire LBD layer rotates as a rigid body 5° perpendicular to the pore axis (**Fig. 4c**).

**Figure 4.**
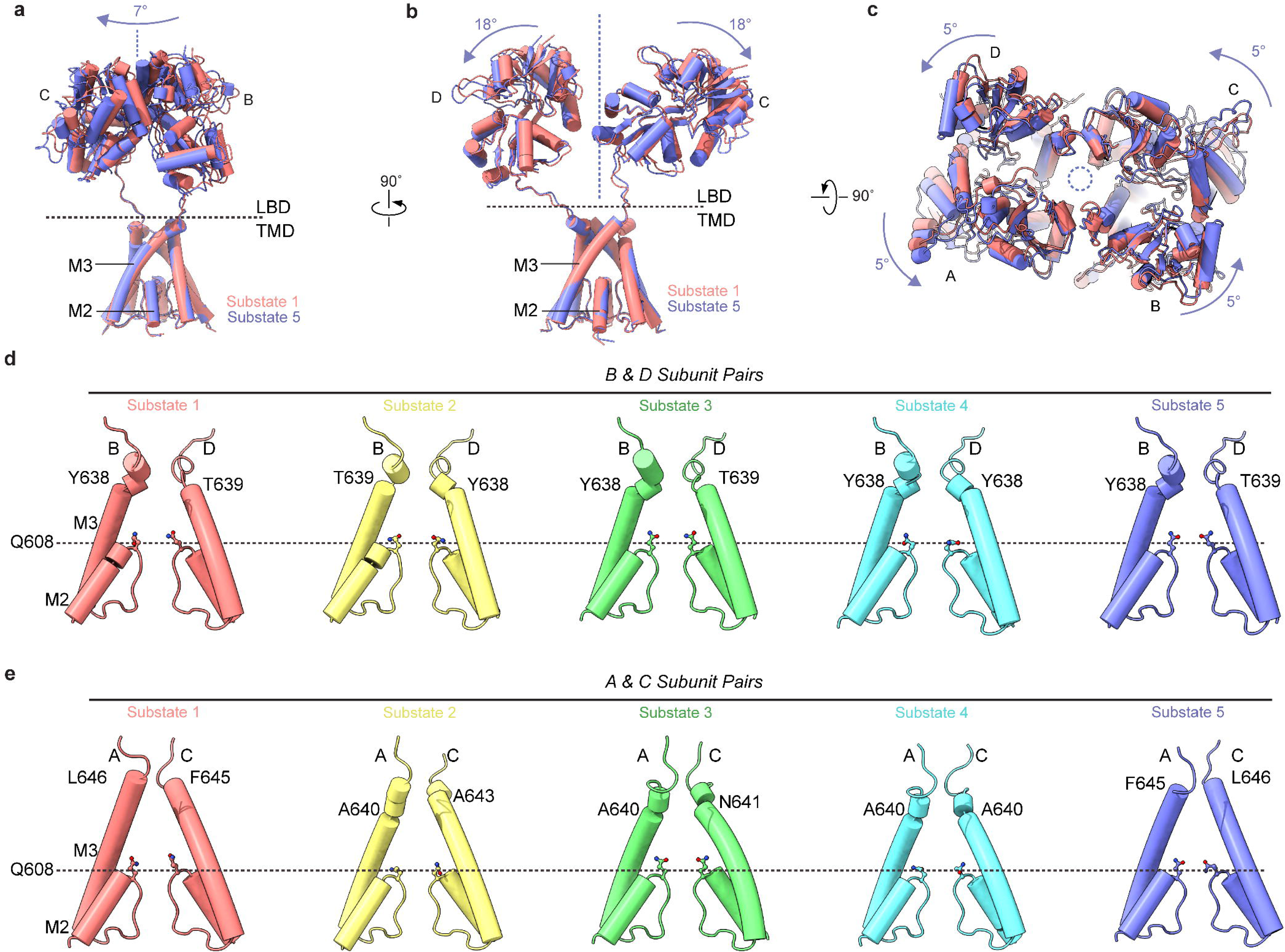
Conformational Plasticity Alters the Gating Hinge. **a** View of the local LBD dimer of substate 1 and substate 5, showing the magnitude of rotation of an individual local dimer. **b** Side view of substate 1 and substate 5, showing 18° of rotation of the LBDs between the two states. **c** Top view of the LBD layer showing 5° of rotation of the LBD layer about the pore axis. **d** B/D helices from each substate showing the location of the gating hinge for each helix. The most prominent gating hinges for the B/D subunits are at Y638 and T639. **e** A/C helices from each substate showing the gating hinge progressing deeper and deeper until reaching A640 in substate 4 and becoming less pronounced in substate 5.

Given that transmission of motion in the LBD to the M3 linkers and TMD underlies channel gating during activation, we investigated how these motions were transferred from the LBD to the TMD layer. Examination of the B and D helices of each substate revealed that the location of the gating hinge changed between substates, hinging at either Y638 or T639 (**Fig. 4d**). Hinging in the A and C helices was more complex with the location of the A and C hinge moving downward through the substates, with the maximally open substate 4 exhibiting the deepest hinge, with both A and C subunits hinging at A640 (**Fig. 4e**). These data indicate that the magnitude of channel opening in GluA4 AMPARs is at least partially controlled by the magnitude and location of the helical kinks in the A and C subunits, and that this can be directly tuned by the conformational plasticity in the LBD layer.

## Discussion

### The GluA4 NTD is intact, but conformationally heterogeneous

We have solved the structure of the disease-relevant AMPAR subunit GluA4 in the active conformation. This structure shows an unprecedented degree of conformational heterogeneity that puts forward a model of AMPARs as dynamic structures with flexible domains that may contribute to or provide resilience to forms of plasticity at synapses. Despite the conformational heterogeneity observed in the GluA4 homotetramer, the overall topology of the receptor is intact with an intact trans-dimer interface between the B and D subunit positions at the level of the NTD

This result is surprising given that recent work has indicated the significance of GluA2 for the overall integrity and topology of AMPARs. GluA2 has been observed preferentially occupying the B/D subunit positions in native receptors^5^, which form the trans-dimer interface in the NTD layer in these receptors as well. In contrast, solution experiments with purified AMPAR NTDs revealed that the NTDs of GluA1 and GluA4 likely exist in a monomer/dimer equilibrium, suggesting that homomeric GluA1 and GluA4 receptors may be less stable at the NTD level^45^. Supporting this idea, a recent study demonstrated that homomeric GluA1 receptors show an NTD that is ruptured at the trans-dimer interface, breaking the classic Y-shape of the receptor. Functional analysis of GluA2 mutated to disrupt the NTD revealed that the disruption of the trans-dimer interface impaired the ability of AMPARs to stably insert at synapses^40^.

Given that the NTD is thought to be important for the biogenesis and assembly of AMPARs, one critical question is how CP-AMPARs, which may lack GluA2, assemble without the GluA2 subunit in the B/D subunit positions. This question is of particular relevance in interneuron classes like PVINs which express GluA4 and GluA1 but little GluA2^9^. The stability of the trans-dimer interface with GluA4 in the B/D position therefore might indicate that in cell types like PVINs, GluA4 may act as a calcium-permeable structural alternative for GluA2, occupying the B/D subunit positions which form the trans-dimer interface and dominating receptor gating. This might suggest that among the four AMPAR subunits GluA1-GluA4, there exists a dichotomy in which GluA2 and GluA4 are competent for assembly at the B/D positions whereas GluA1 and GluA3 are not. Given the dominance of the B and D subunits for gating of the receptor, this dichotomy would suggest that GluA2 and GluA4 exhibit outsize influence on the physiological properties of the receptor.

### Activation of GluA4

The conformational heterogeneity in the GluA4 NTD layer supports previous notions that this domain of the receptor may serve as a platform for interactions with other proteins, but the heterogeneity of this layer did not prevent the protein from entering the active state in the presence of 1mM Glu and 100μM CTZ (**Fig 2).** This active state is characterized by kinking at all four helices, similar to what has been recently observed with GluA2 AMPARs activated at physiological temperatures^43^. However, activation of GluA4 exhibits kinking of the M3 helices at the previously unappreciated residues Y638 and T639 in the B/D helices and L642 in the A/C helices. In contrast to the previously solved active states of GluA2, the GluA4 active state is maximally constricted at T639 rather than at the more common A643 constriction. This difference in gating mechanism may explain differences in GluA4 gating kinetics such as rapid desensitization. Together, T639 and A640 are homologous to the calcium-binding ‘Site G’ recently described in GluA2 homomers^8^.

### Conformational Plasticity in GluA4 Tunes the Channel Gate

Through analyzing the variability in our GluA4 active state consensus dataset, we were able to identify that the GluA4 active state was not a single state but an average along a continuum of heterogeneity in the position of the LBD relative to the TMD. The variability along this axis not only changed the relative position of the LBD to the TMD, but also altered the radius of the channel pore at the upper channel gate by kinking the M3 helices of the A and C subunits deeper and deeper within the pore, achieving the most open state (Substate 4) by kinking the A/C helices at A640. Among all substates, we have identified eight residues which can kink in the M3 helices to facilitate channel gating (Y638, T639, A640, N641, L642, A643, F645, L646). Of these only A640 (A618 in GluA2, the “A-hinge”)^35,43^ has been previously reported. When including the “S-hinge” and the “Lurcher hinge” recently identified in glutamate-gated AMPARs among the list of hinging residues^43^, it appears that every residue in the 637-SYTANLAAF-645 motif common to ionotropic glutamate receptors is competent for forming a gating hinge, explaining why this motif is so thoroughly conserved among ionotropic glutamate receptors^1^ and why mutations in this motif produce pathological outcomes in human patients. Indeed, T639, N641, and A643 have recently been identified in human patients experiencing intellectual disability and epilepsy^16^.

AMPARs are known to open to four different subconductance states (O1-O4) although the mechanism of subconductance states and how they are achieved has been the subject of debate. Longstanding models in the field have posited that subconductance states are achieved through differing occupancies of the glutamate-binding LBDs and that conductance scales with the number of LBDs bound to the neurotransmitter glutamate^39,46–54^. However, a recent study examining the magnitude of channel openings in AMPARs with variable glutamates bound has called this view into question. When subsaturating glutamate concentrations were applied to purified AMPARs in the presence of the desensitization-blocking PAM CTZ, a mixed population of subunit occupancies, with receptors bound to 0,1,2,3 or 4 glutamate molecules were observed via cryoEM^39^. In these conditions, the magnitude of channel opening did not correlate with the number of glutamates bound, with the highest channel openings achieved through 2 glutamates bound and 3 and 4 glutamate-bound AMPARs showing a maximal channel opening of 1.7 Å. These findings suggest that the subunit occupancy model of subconductance may be oversimplified. We show here that a fully saturated receptor produces channel openings comparable to those and even exceeding the range observed with subsaturating glutamate^39^. We propose that conformational heterogeneity is an alternative mechanism by which subconductance may be achieved in AMPARs. We suspect that these two mechanisms may converge on a common structural mechanism of subconductance, perhaps having to do with kinetics of clamshell closure following ligand binding to the AMPAR^54^.

## Methods

### Construct Design

A fusion construct consisting of the coding sequence of GluA4 fused to TARPγ2 was generated based on previous strategies for generating AMPAR expression constructs and a similar construct used for studying GluA4 physiology (**Extended Data 1**)^33–35,37,38,40^. This construct is referred to as GluA4-γ2 and was prepared as follows. Rat GluA4 was truncated after the fourth transmembrane helix and a single Glycine-Threonine (GT) linker was inserted between the GluA4 truncation and the N-terminus of the human TARPγ2 coding sequence. The TARPγ2 coding sequence was itself truncated after the fourth transmembrane helix (TM4), followed by a Threonine-Glycine-Glycine (TGG) linker, a Thrombin cleavage site, a short polyalanine stretch and then the coding sequence for eGFP. Finally, a Strep Tag ii was appended to the end of the coding sequence prior to the stop codon to facilitate protein purification.

### Protein Expression and Purification

Bacmid for the expression of GluA4-γ2 was prepared as previously described^34–37^. P1 baculovirus was generated by transfecting ExpiSf9 cells (Gibco, A35243) maintained at 27 °C with polyethyleneimine (PEI, molecular weight 40,000; PolyScience, 24765). After 5 days P1 baculovirus was harvested and used to induce expression in mammalian cells. Media containing P1 virus was added to cultures of Expi293F GnTI^-^ cells (Gibco, A39240) grown to a density of 2.5 x 10^6^ cells/mL in Expi293 medium (Gibco, A14135101) at a ratio of 1:10 P1 virus to Expi293F culture volume. Infected Expi293F cells were incubated at 37 °C for 12-18 hours before bringing the cuture medium up to 10mM sodium butyrate (Sigma, 303410) and 2μM ZK 20075 (Tocris, 2345) and incubating the culture at 30 °C in 5% CO_2_. 72 hours after transduction, the cells were harvested by low-speed centrifugation (5,000xg, 20 min at 4 °C), washed with PBS (pH 7.4) with protease inhibitors (0.8 μM aprotinin, 2 μg/ml leupeptin, 2 μm pepstatin A and 1 mM phenylmethylsulfonyl fluoride) and then centrifuged a second time (4,800xg, 10 min at 4 °C). Supernatant from this spin was discarded and the cell pellets were stored at -80 °C until purification. On the day of purification, pellets were thawed by rotating the pellet in chilled lysis buffer (150 mM NaCl, 20mM Tris pH 8.0) with protease inhibitors added. After thawing, cells were lysed in an ice bath with a blunt probe sonicator (three cycles of 1s on, 1s off for 1 min, 20W power), and the sonicated lysate was gently centrifuged (4,800xg, 20 min at 4 °C) to clear large cellular debris. The supernatant was ultracentrifuged (125,000xg, 45 min at 4 °C) to pellet membranes. Solubilization buffer (150mM NaCl, 20mM Tris pH 8.0, 1% *n*-dodecyl-β-D-maltopyranoside (DDM; Anatrace D310) and 0.2% Porcine Brain Total Lipid Extract (Avanti, 131101P)) was added to the pelleted membranes and membranes were solubilized for 2 hours at 4 °C in a cold room under constant agitation. Solubilized membranes were then ultracentrifuged (125,000xg, 45 min at 4 °C) to remove insoluble material and the supernatant was applied to a pre-equilibrated volume of Strep-tactin XT 4Flow resin (IBA, 2-5010) overnight, rotating at 4 °C. The next day, the resin was collected by gravity flow and washed with 20 volumes of glycol-diosgenin (GDN) buffer (150 mM NaCl, 20mM Tris pH 8.0 and 0.01% GDN (Anatrace, GDN101)). Bound protein was eluted with 50 mM D-biotin in GDN buffer and collected directly into a centrifugal concentrator and concentrated into a 500 μl volume at 4 °C. The concentrated protein was then treated with thrombin at a 1:200 w/w ratio for 1 h at 22 °C to cleave off eGFP and Strep Tag II. The cleavage reaction was separated over a Superose 6 increase 10/300 column (Cytiva, 29091596) using an AKTA fast protein liquid chromatograph in GDN buffer. Two peaks were observed on the chromatogram corresponding to a higher and lower molecular weight species for GluA4. Peak fractions corresponding to the lower molecular weight species were collected and concentrated in a centrifugal concentrator to 1.49 mg/ml.

### Cryo-EM sample preparation and data collection

Purified GluA4/TARPγ2 was treated with 100μM cyclothiazide (CTZ, Tocris, 07-131-0) and ultracentrifuged (125,000xg, 45 min at 4 °C) to remove precipitated protein prior to grid preparation. Au-Flat 1.2 μm/1.3 μm 300 mesh gold grids (MiTeGen, M-CEM-AUFT313) were plasma treated in a Pelco EasyGlow (25 mA, 120s glow time and 10s hold time; Ted Pella, 91000). The GluA4/TARPγ2 complexed with CTZ was spiked with glutamate (pH 7.4) immediately prior to the preparation of grids. To prepare grids, 3 μl of sample was applied to glow-discharged grids in an FEI Vitrobot Mark IV (Thermo Fisher Scientific; wait time 30s; blot force 5, blot time 4 s) set to 8 °C and 100% humidity and plunge frozen in liquid ethane. Grids were imaged with a 300-kV Titan Krios 3i microscope equipped with fringe-free imaging, a Falcon 4i camera and a Selectris energy filter set to a 10-eV slit width. Micrographs were collected with a dose rate of 3.83 e^-^ per pixel per s and a total dose of 40.00 e^-^ per Å^2^. We collected 19,054 micrographs with a pixel size of 0.93 Å per pixel. Automated collection was achieved with EPU software from ThermoFisher Scientific.

### Image processing

Cryosparc^55^ was used for all aspects of image processing (refer to Extended Data Figs. 2 and 7 for details). The reconstruction quality was tested for anisotropic contribution to the Fourier shell correlation (FSC) with 3DFSC.

### Model building, refinement, and structural analysis

Molecular modeling was performed in ChimeraX^56^ and Coot^57^. Refinement was performed with a combination of ISOLDE^58^ and PHENIX^59^. These software packages were accessed through the SBgrid consortium^60^. As a starting model, four copies of the LBD and TMD layers of AlphaFold 2 prediction for GluA4 were each rigid-body fit into the consensus model and then each protomer LBD and TMD layer was joined. Four copies of the previously solved model of TARPγ2 was then rigid body fit into each corresponding density to create a single consensus model. ISOLDE was then used to adjust the position of all amino acids simultaneously before inspecting each amino acid fit to the map and merging Glu and CTZ into the model in Coot. Phenix was used to refine the final model. MolProbity^61^ was used to assess model quality. Visualization, figure generation and domain measurements were performed in ChimeraX. HOLE was used to measure the radius of the ion channel pore along the z axis^44^.

### Single channel recording

Single GluA4/TARPγ2 channels were recorded in reconstituted lipid bilayers using Meca 4 Recording chips (100μm cavity; Nanion, 132002) in an Orbit Mini device (Nanion) using the Elements Data Reader software (Elements S.R.L./Nanion) as previously reported^39^. Briefly, 1,2-diphytanoyl-sn-glycero-3-phosphocholine (DPHPC; Avanti Polar Lipids 850356P25MG) was solubilized at 10mg/ml in decane (Sigma, 803405) and used to paint lipid bilayers onto a recording chip equilibrated in recording solution (150mM KCl, 20 mM HEPES pH 7.2, 1mM Glutamate, 100 μM CTZ). GluA4/TARPγ2 sample was diluted to 10-100 ng/ml and mixed 1:1 with DPHPC solution and placed on a heat block at 37 °C for 30 min. Each recording channel was repainted with protein/lipid mixture until each aperture displayed a capacitance between 5 and 15 pF. Recordings were sampled at 20kHz and filtered at 1.2kHz. Analysis of recordings was performed in Clampfit 11 (Axon Devices) following the application of an 8-pole Bessel filter to smooth the recordings. Analysis was restricted to 500 ms epochs surrounding ‘high P_open_’ events.

## Data Availability

The structural coordinates and maps will be available upon publication. Source data are provided with this paper.

## Acknowledgements

We would like to thank R. Johnson and C. Warrick for their assistance with developing the GluA4 expression system. We also thank members of the Huganir and Twomey labs for comments and helpful discussion. All cryo-EM data were collected at the Beckman Center for Cryo-EM at Johns Hopkins with assistance from D. Sousa, D. Ding and K. Cai. W.D. H is supported by National Institutes of Health (NIH) grant K99 MH132811. R.L.H. is supported by NIH grant R37 NS036715. E.C.T. is supported by NIH grant R35GM154904, the Searle Scholars Program (Kinship Foundation 22098168) and the Diana Helis Henry Medical Research Foundation (142548).

## Author Contributions

W.D.H., R.L.H. and E.C.T. designed the study. W.D.H carried out all experiments. W.D.H., R.L.H. and E.C.T. analyzed the data and wrote the paper. R.L.H. and E.C.T. supervised the project.

## Extended Data Figure Legend

**Extended Data 1.**
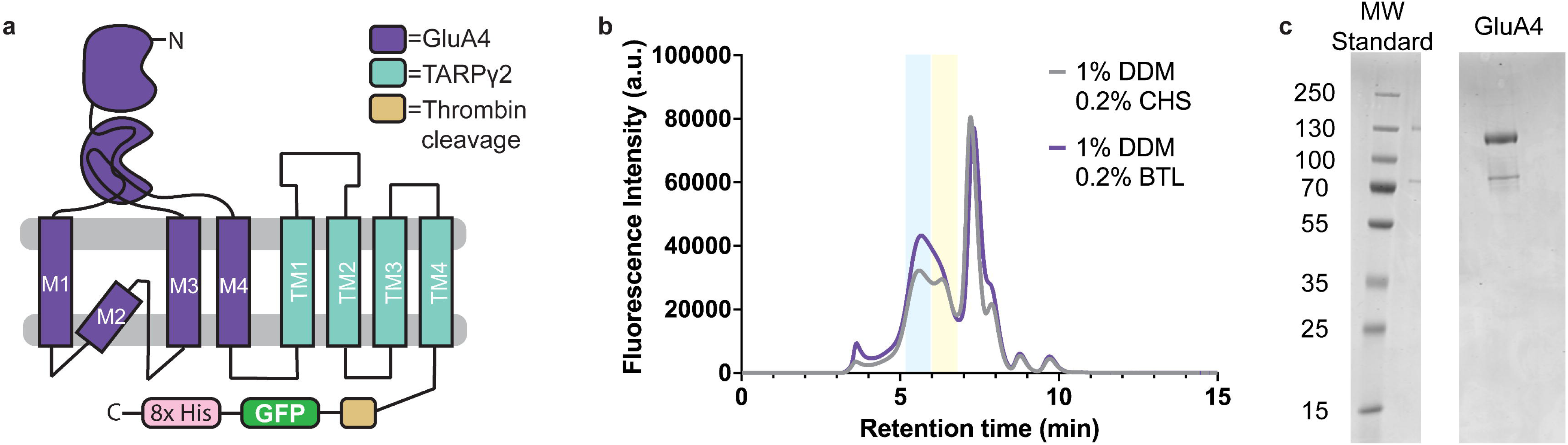
GluA4/TARPγ2 purification strategy. **a** GluA4/TARPγ2 construct strategy showing the fusion between GluA4 (purple), which is concatemerized with TARPγ2 (teal) by truncating the GluA4 coding sequence following the fourth transmembrane domain (M4) and fusing this directly to the N-terminus of TARPγ2. **b** Fluorescence size-exclusion chromatogram of GluA4/TARPγ2 protein. The gray trace is from protein solubilized in buffer containing 1% DDM, 0.2% cholesteryl hemisuccinate (CHS), while the purple trace is from protein solubilized in 1% DDM, 0.2% porcine brain total lipid extract (BTL). The vertical light blue bar represents the predicted size of a tetrameric AMPAR fully occupied by TARPγ2 while the tan bar represents the predicted size of a single GluA4/TARPγ2 monomer. **c** Coomassie stained SDS PAGE gel of purified GluA4/TARPγ2 showing the prominent band at the expected molecular weight of ∼120kD.

**Extended Data 2.**
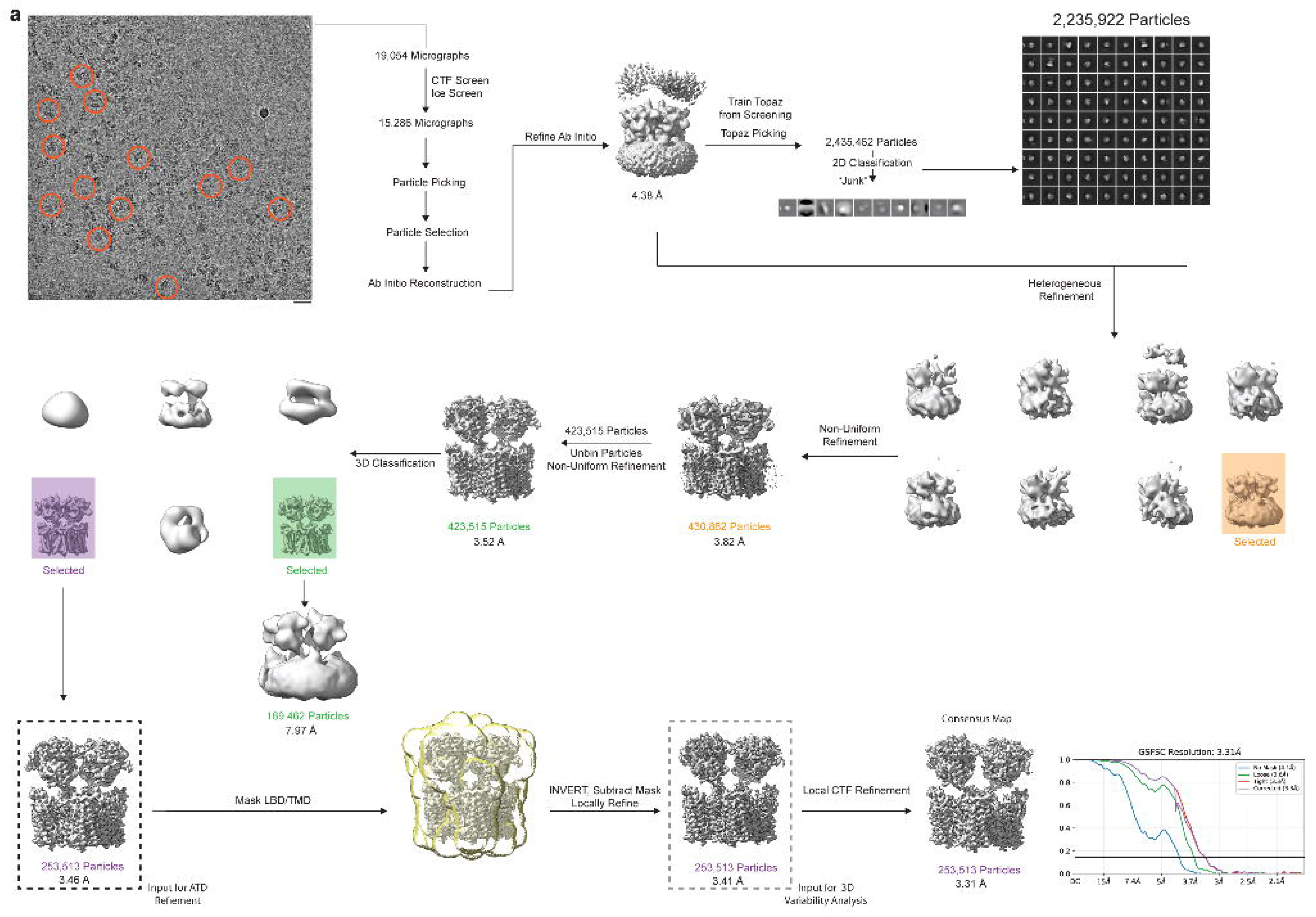
Workflow for processing GluA4/TARPγ2 active state micrographs. Diagram showing the overall workflow for processing the GluA4/TARPγ2 data from micrographs to the consensus LBD/TMD assembly (Consensus Map). The black dashed box highlights the input for NTD refinement (Extended Data 3) whereas the gray dashed box indicates the input for 3D Variability Analysis of the LBD/TMD assembly (Extended Data 7).

**Extended Data 3.**
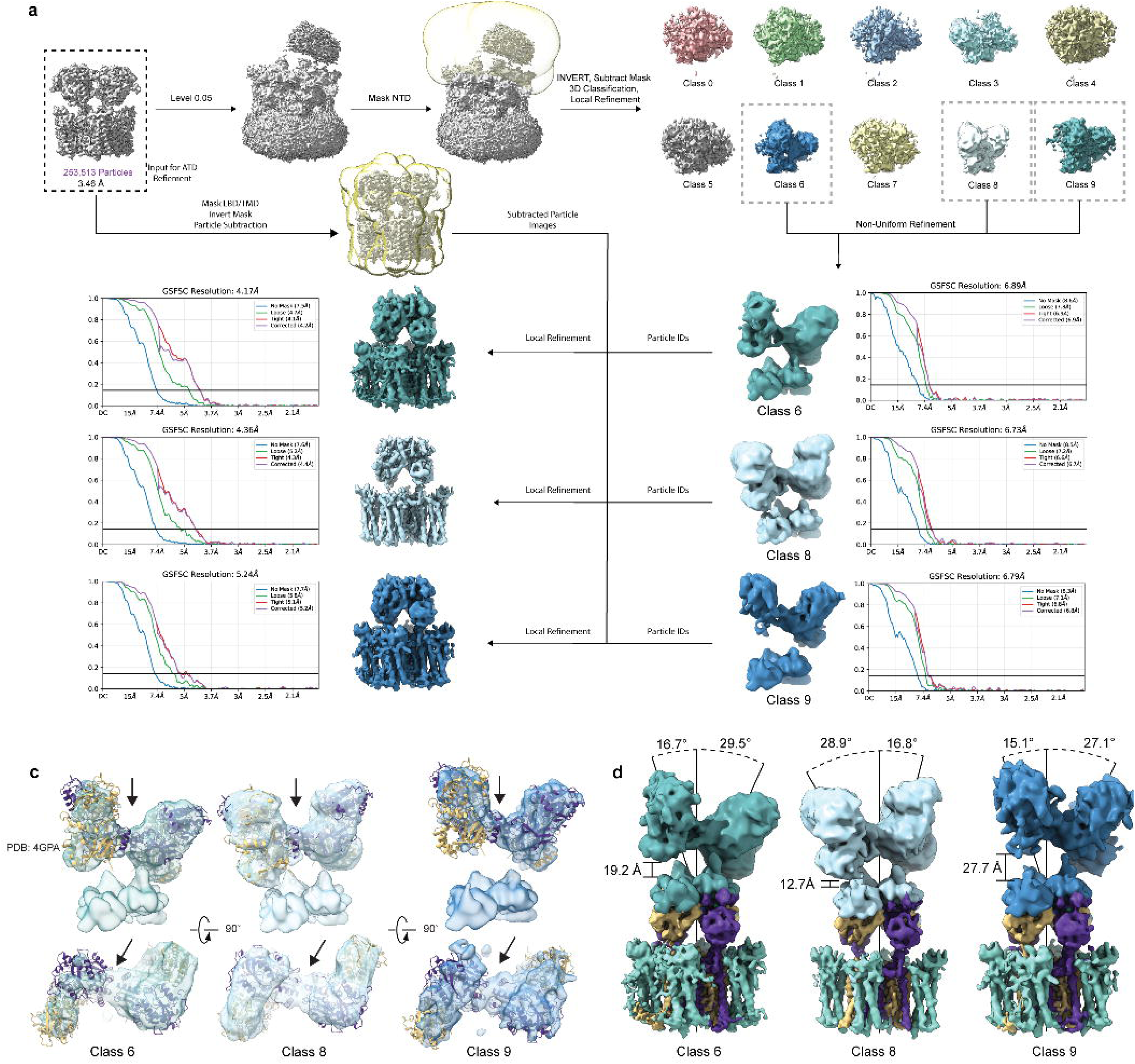
Strategy for NTD refinement and Analysis of NTD geometry. **a** Workflow for refining the NTD density from the intact GluA4/TARPγ2 map (black dashed box). **b** Analysis of the three resolvable NTD classes showing the fit of the existing GluA4 NTD crystal structure within the density (PDB: 4GPA). NTD models are colored according to their subunit identity in the overall reconstruction. The trans-dimer interface is shown with an arrow. **c** Composite maps created by aligning the NTD alone and LBD/TMD maps (from **a**). The position of the NTD relative to the LBD/TMD assembly was measured using models fit into the respective densities in ChimeraX.

**Extended Data 4.**
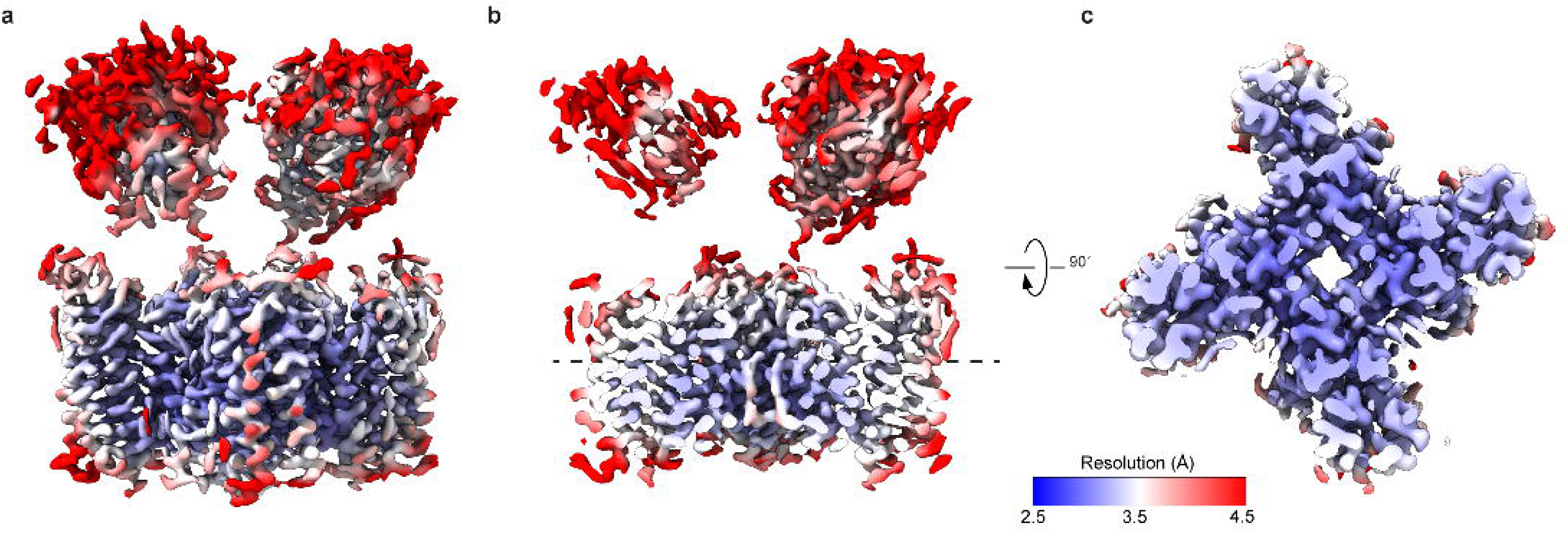
Local resolution of the GluA4/TARPγ2 consensus map. **a** Local resolution mapped onto the overall GluA4/TARPγ2 structure. **b** Coronal section through the GluA4/TARPγ2 map showing the inside of the channel pore as well as the middle of the LBD lobes. **c** Section through the transmembrane domain at the level of the dashed line in **b**, showing the TARP and GluA4 TM helices surrounding the pore. Resolution is mapped between 2.5 Å (dark blue) and 4.5 Å (red).

**Extended Data 5.**
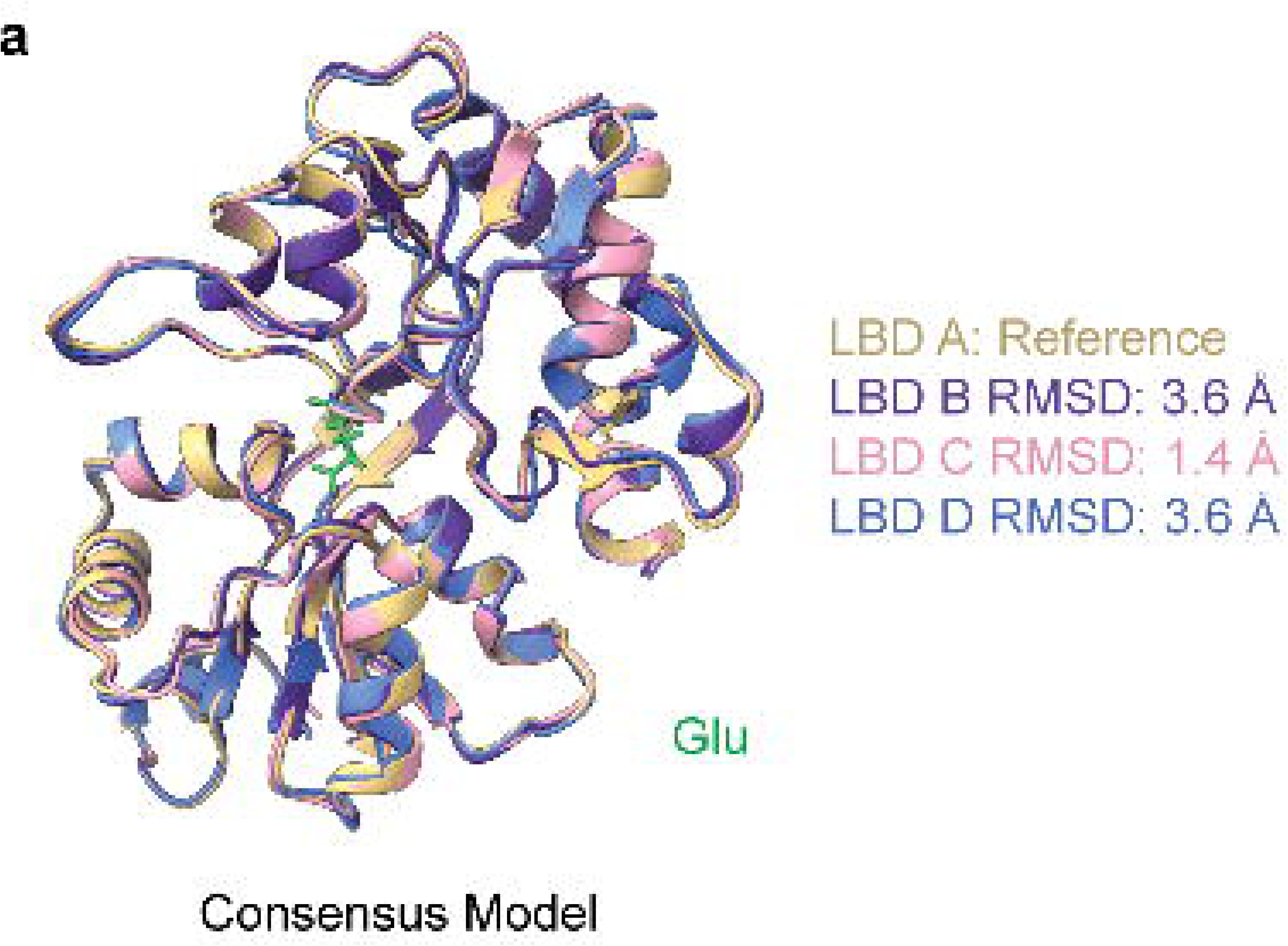
All four GluA4/TARPγ2 LBDs are bound to Glu. Overlay of the isolated LBDs from the GluA4/TARPγ2 model (tan, purple, blue, pink) showing that each is similarly closed around Glu (green) bound in the LBD clamshell cleft.

**Extended Data 6.**
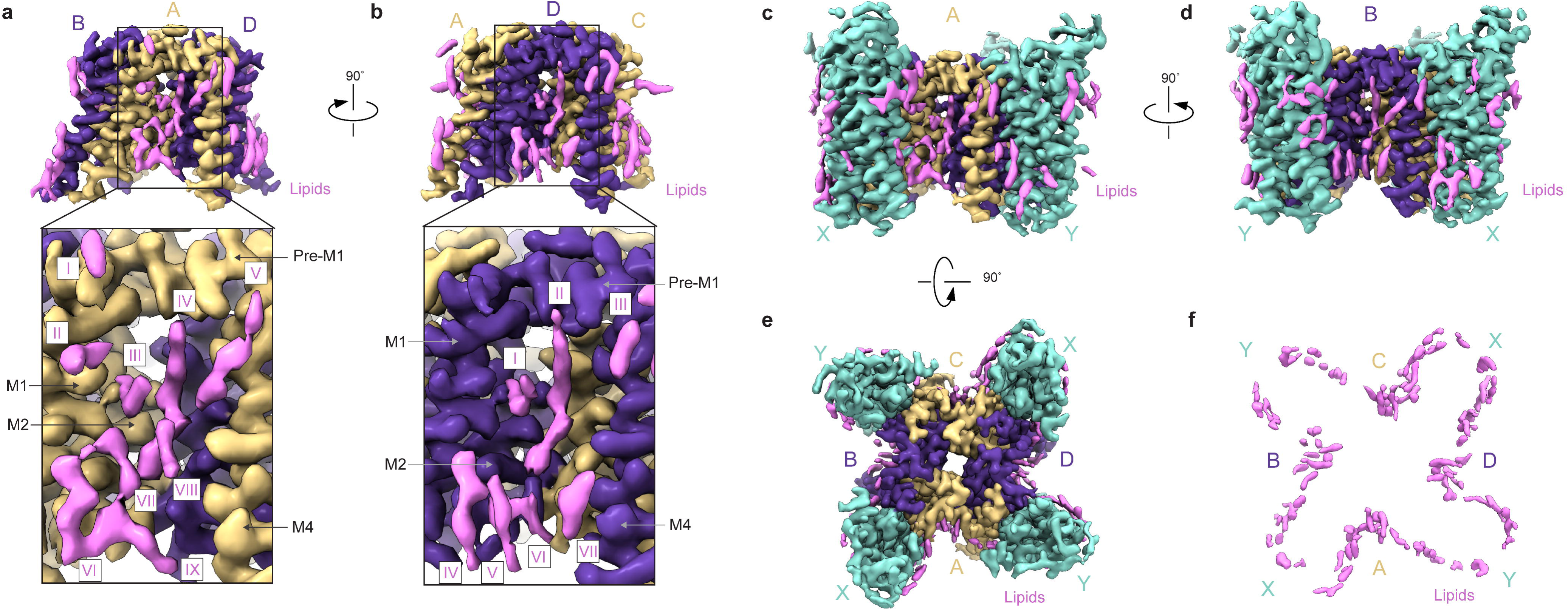
Lipids bound to the GluA4-TARPγ2 TMD. **a** Representative fenestral lipids bound to the A/C subunit positions of GluA4. Nine overall chains decorate the fenestra of the A/C subunits, with lipids II and III angled towards the vestibule of the pore. **b** Representative fenestral lipids bound to the B/D subunit positions of GluA4. Fewer distinct lipids are observable at this position. Lipid I is angled towards the pore vestibule like lipid III in a. **c** Lipid decoration of the GluA4-TARPγ2 complex, with the A subunit position centered. Compared to a, additional lipid residues can be seen forming the interface between GluA4 and TARPγ2 as well as lipids that bind solely to TARPγ2. **d** Similar to c but showing lipids visible with the B subunit of GluA4 visible. **e** Top-down view of the GluA4 pore showing lipids in pink surrounding the GluA4 TMD assembly. **f** Bound lipids displayed alone, revealing the lipid footprint of the GluA4-TARPγ2 complex.

**Extended Data 7.**
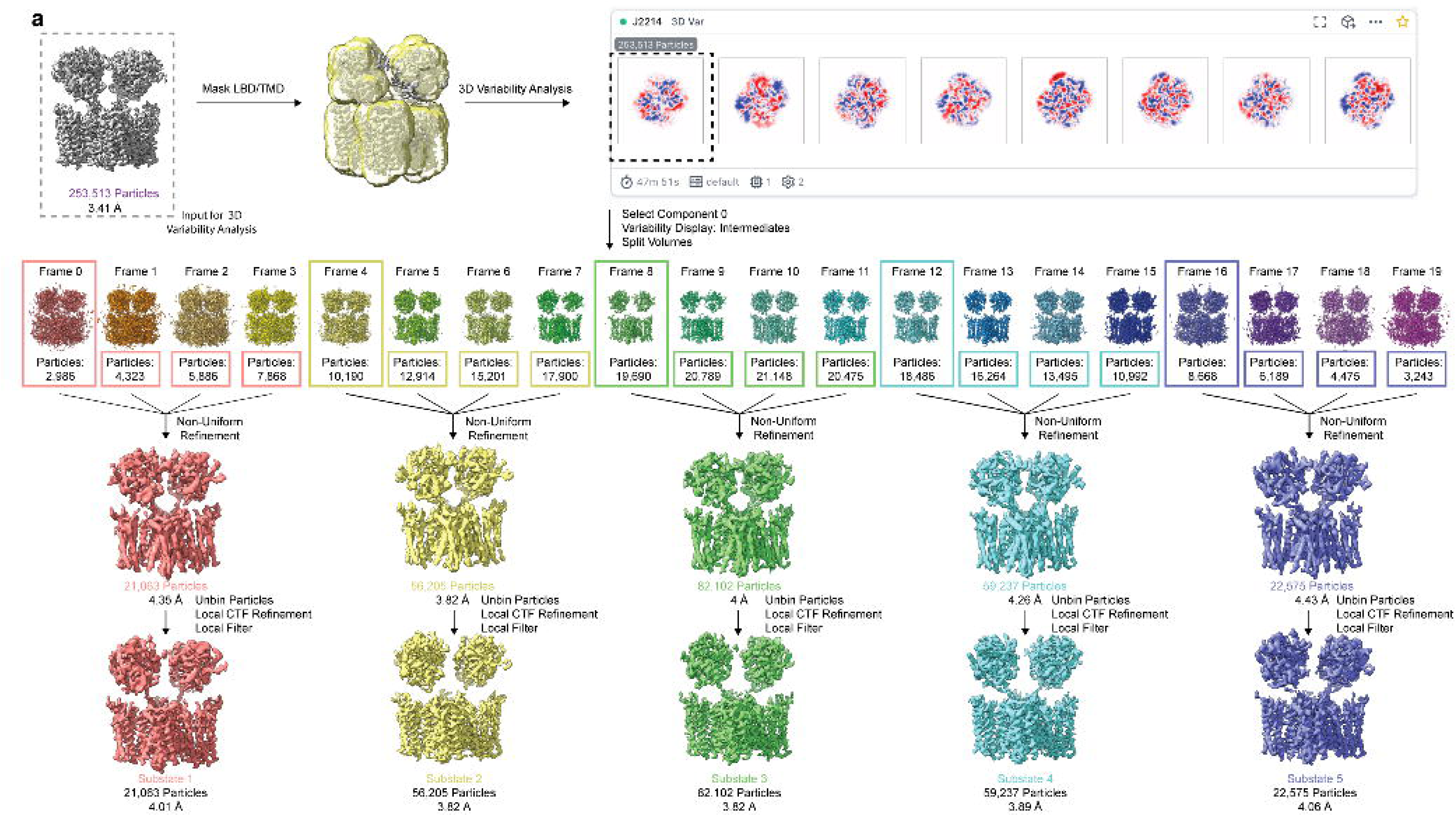
Workflow for 3D Variability Analysis of the consensus map. Overall workflow of 3D Variability Analysis of the LBD/TMD maps of the GluA4/TARPγ2 complex. The unfiltered LBD/TMD map from Extended Data 2 (gray dashed box) was used as input for 3D Variability Analysis.

**Extended Data 8.**
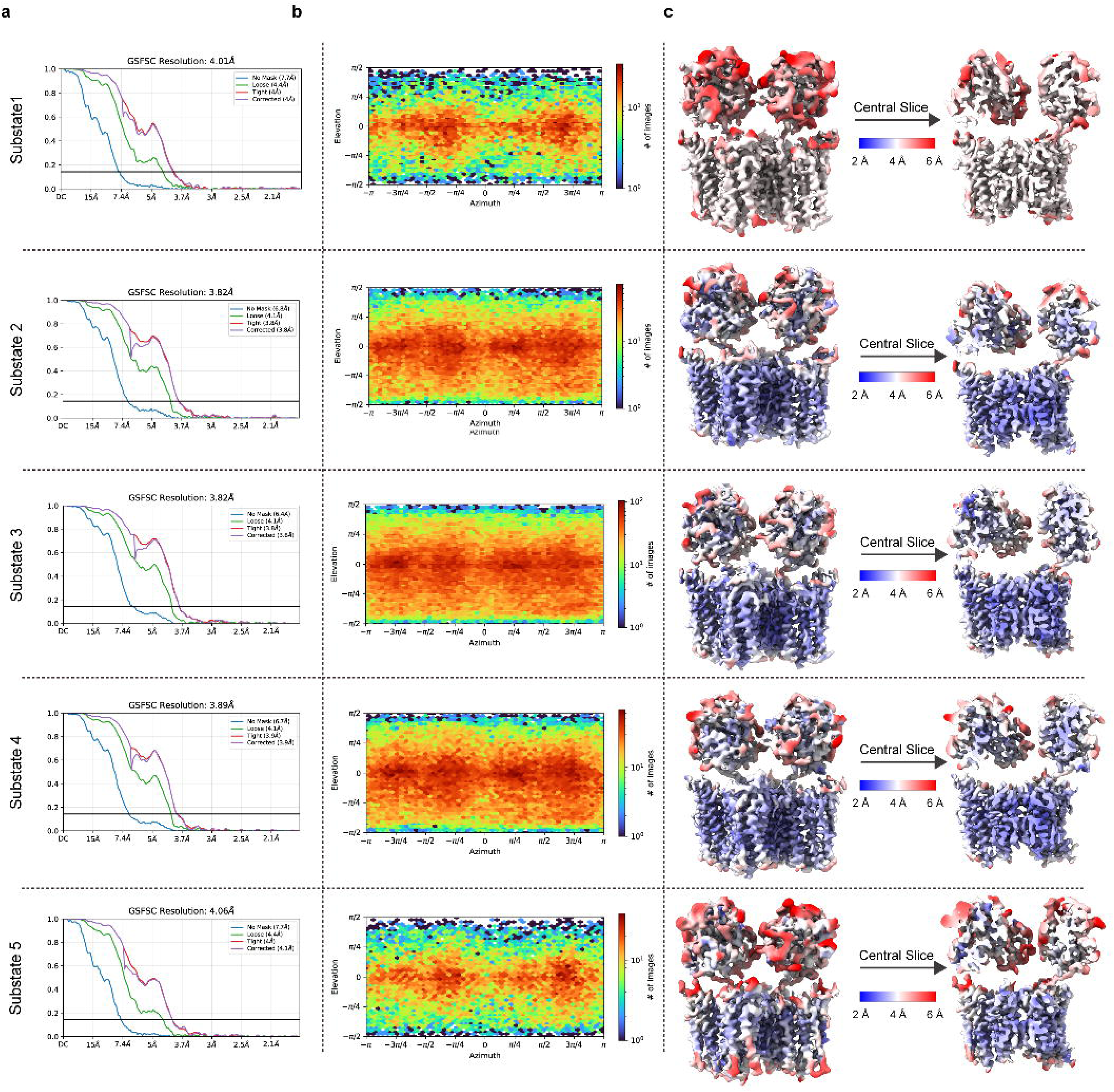
Local resolution maps of GluA4 substates. **a** GSFC curves corresponding to each substate. **b** Euler angle plots for each GluA4 substate. **c** Local resolution maps for each GluA4 substate.

**Extended Data 9.**
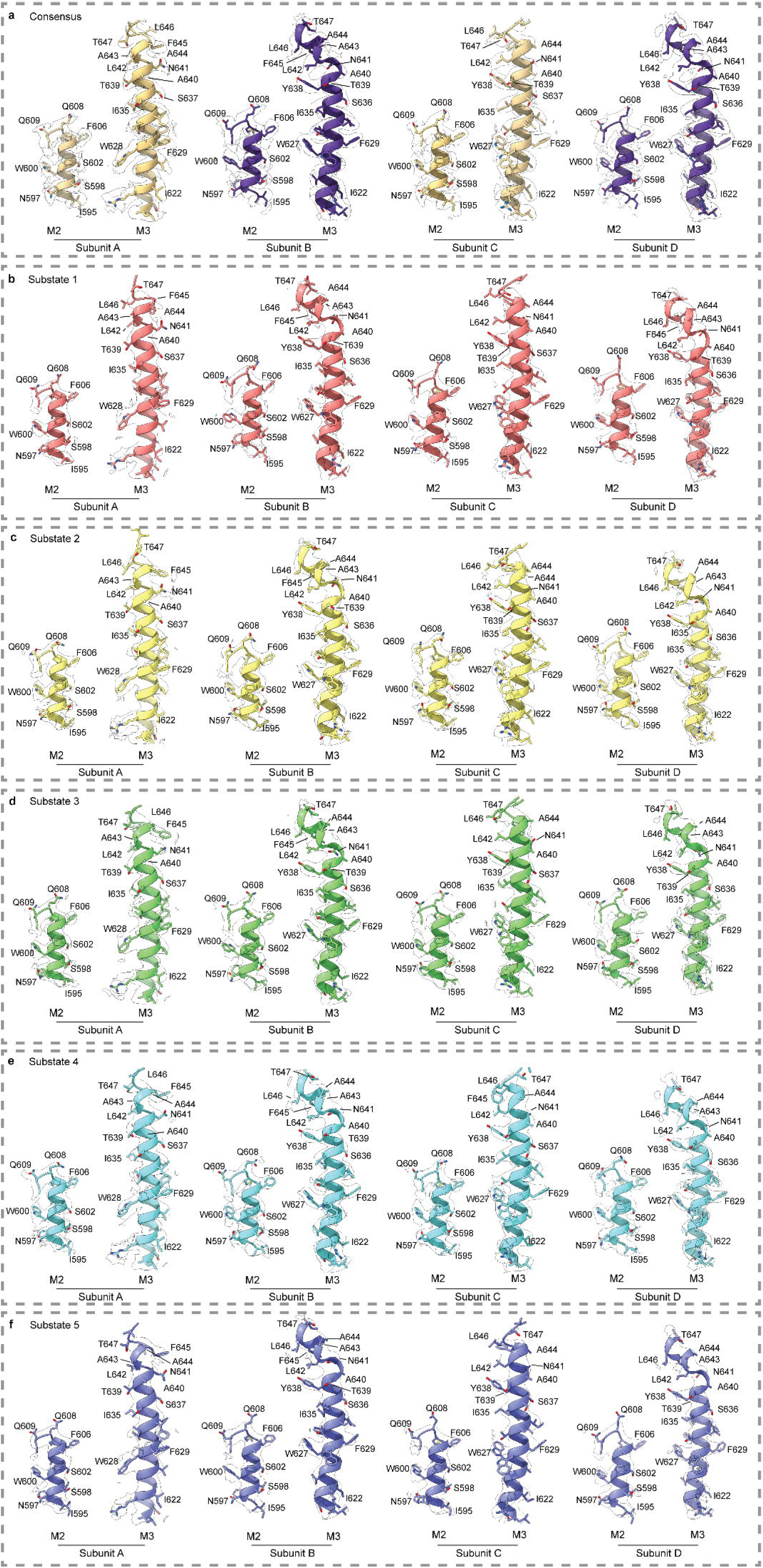
Map fit of the core pore-forming helices among GluA4 substates. a-f. Map versus model fit of the consensus state pore-forming helices (**a**) and the substate pore forming helices (**b-f**). The map is shown as a black outline around each model and is contoured at 0.14 for each substate.

**Extended Data 10.**
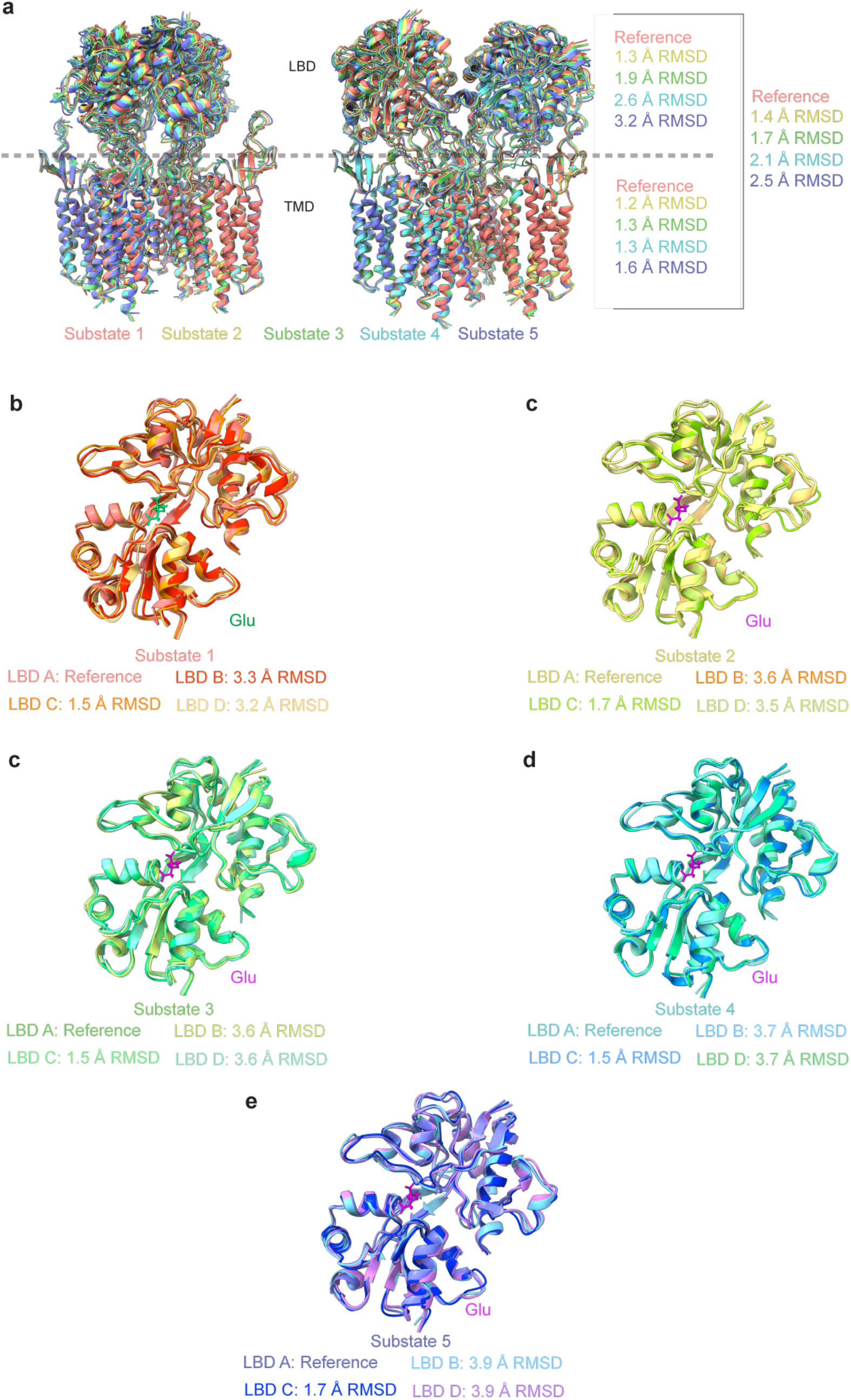
Comparison of substate models. **a** Alignment of the substate models show high agreement in the TMD, but conformational heterogeneity in the position of the LBD relative to the TMD. **b-e** Overaly of the LBDs from all four subunit positions (A-D) for each substate (1-5). Glu is colored green in **b** and fuchsia in **c-e**.

## References

1. Hansen, K. B. et al. Structure, Function, and Pharmacology of Glutamate Receptor Ion Channels. Pharmacol. Rev. 73, 1469–1658 (2021).

2. Twomey, E. C. & Sobolevsky, A. I. Structural Mechanisms of Gating in Ionotropic Glutamate Receptors. Biochemistry 57, 267–276 (2018).

3. Keinänen, K. et al. A Family of AMPA-Selective Glutamate Receptors. Science 249, 556– 560 (1990).

4. Diering, G. H. & Huganir, R. L. The AMPA Receptor Code of Synaptic Plasticity. Neuron 100, 314–329 (2018).

5. Zhao, Y., Chen, S., Swensen, A. C., Qian, W.-J. & Gouaux, E. Architecture and subunit arrangement of native AMPA receptors elucidated by cryo-EM. Science 364, 355–362 (2019).

6. Cull-Candy, S. G. & Farrant, M. Ca2+-permeable AMPA receptors and their auxiliary subunits in synaptic plasticity and disease. J. Physiol. 599, 2655–2671 (2021).

7. Miguez-Cabello, F. et al. GluA2-containing AMPA receptors form a continuum of Ca2+-permeable channels. Nature 641, 537–544 (2025).

8. Nakagawa, T., Wang, X., Miguez-Cabello, F. J. & Bowie, D. The open gate of the AMPA receptor forms a Ca2+ binding site critical in regulating ion transport. Nat. Struct. Mol. Biol. 31, 688–700 (2024).

9. Hong, I. et al. Calcium-permeable AMPA receptors govern PV neuron feature selectivity. Nature 635, 398–405 (2024).

10. Kwok, K. H. H., Tse, Y. C., Wong, R. N. S. & Yung, K. K. L. Cellular localization of GluR1, GluR2/3 and GluR4 glutamate receptor subunits in neurons of the rat neostriatum. Brain Res. 778, 43–55 (1997).

11. Mosbacher, J. et al. A Molecular Determinant for Submillisecond Desensitization in Glutamate Receptors. Science 266, 1059–1062 (1994).

12. Geiger, J. R. P. et al. Relative abundance of subunit mRNAs determines gating and Ca2+ permeability of AMPA receptors in principal neurons and interneurons in rat CNS. Neuron 15, 193–204 (1995).

13. Swanson, G. T., Kamboj, S. K. & Cull-Candy, S. G. Single-Channel Properties of Recombinant AMPA Receptors Depend on RNA Editing, Splice Variation, and Subunit Composition. J. Neurosci. 17, 58–69 (1997).

14. Chang, M. C. et al. Narp regulates homeostatic scaling of excitatory synapses on parvalbumin-expressing interneurons. Nat. Neurosci. 13, 1090–1097 (2010).

15. Pelkey, K. A. et al. Pentraxins Coordinate Excitatory Synapse Maturation and Circuit Integration of Parvalbumin Interneurons. Neuron 85, 1257–1272 (2015).

16. XiangWei, W. et al. Clinical and functional consequences of GRIA variants in patients with neurological diseases. Cell. Mol. Life Sci. 80, 345 (2023).

17. Wang, Y.-X., Wenthold, R. J., Ottersen, O. P. & Petralia, R. S. Endbulb Synapses in the Anteroventral Cochlear Nucleus Express a Specific Subset of AMPA-Type Glutamate Receptor Subunits. J. Neurosci. 18, 1148–1160 (1998).

18. Gardner, S. M., Trussell, L. O. & Oertel, D. Time Course and Permeation of Synaptic AMPA Receptors in Cochlear Nuclear Neurons Correlate with Input. J. Neurosci. 19, 8721–8729 (1999).

19. Rubio, M. E. & Wenthold, R. J. Glutamate Receptors Are Selectively Targeted to Postsynaptic Sites in Neurons. Neuron 18, 939–950 (1997).

20. Hunter, C., Petralia, R. S., Vu, T. & Wenthold, R. J. Expression of AMPA-selective glutamate receptor subunits in morphologically defined neurons of the mammalian cochlear nucleus. J. Neurosci. 13, 1932–1946 (1993).

21. Schmid, S., Guthmann, A., Ruppersberg, J. P. & Herbert, H. Expression of AMPA receptor subunit flip/flop splice variants in the rat auditory brainstem and inferior colliculus. J. Comp. Neurol. 430, 160–171 (2001).

22. Rubio, M. E. et al. The number and distribution of AMPA receptor channels containing fast kinetic GluA3 and GluA4 subunits at auditory nerve synapses depend on the target cells. Brain Struct. Funct. 222, 3375–3393 (2017).

23. Schwenk, J. et al. Regional Diversity and Developmental Dynamics of the AMPA-Receptor Proteome in the Mammalian Brain. Neuron 84, 41–54 (2014).

24. Gonzalez-Lozano, M. A. et al. Stitching the synapse: Cross-linking mass spectrometry into resolving synaptic protein interactions. Sci. Adv. 6, eaax5783 (2020).

25. Kita, K. et al. GluA4 facilitates cerebellar expansion coding and enables associative memory formation. eLife https://elifesciences.org/articles/65152 (2021) doi:10.7554/eLife.65152.

26. García-Hernández, S. & Rubio, M. E. Role of GluA4 in the acoustic and tactile startle responses. Hear. Res. 414, 108410 (2022).

27. Yang, Y.-M. et al. GluA4 is indispensable for driving fast neurotransmission across a high-fidelity central synapse. J. Physiol. 589, 4209–4227 (2011).

28. Taylor, K. R. et al. Glioma synapses recruit mechanisms of adaptive plasticity. Nature 623, 366–374 (2023).

29. Stepulak, A. et al. Expression of glutamate receptor subunits in human cancers. Histochem. Cell Biol. 132, 435–445 (2009).

30. Brocke, K. S. et al. Glutamate receptors in pediatric tumors of the central nervous system. Cancer Biol. Ther. 9, 455–468 (2010).

31. Tao, Y. et al. Erbin interacts with TARP γ-2 for surface expression of AMPA receptors in cortical interneurons. Nat. Neurosci. 16, 290–299 (2013).

32. Tomita, S. et al. Functional studies and distribution define a family of transmembrane AMPA receptor regulatory proteins. J. Cell Biol. 161, 805–816 (2003).

33. Zhang, W., Devi, S. P. S., Tomita, S. & Howe, J. R. Auxiliary proteins promote modal gating of AMPA-and kainate-type glutamate receptors. Eur. J. Neurosci. 39, 1138–1147 (2014).

34. Twomey, E. C., Yelshanskaya, M. V., Grassucci, R. A., Frank, J. & Sobolevsky, A. I. Elucidation of AMPA receptor-stargazin complexes by cryo-electron microscopy. Science 353, 83–86 (2016).

35. Twomey, E. C., Yelshanskaya, M. V., Grassucci, R. A., Frank, J. & Sobolevsky, A. I. Channel opening and gating mechanism in AMPA-subtype glutamate receptors. Nature 549, 60–65 (2017).

36. Twomey, E. C., Yelshanskaya, M. V., Grassucci, R. A., Frank, J. & Sobolevsky, A. I. Structural Bases of Desensitization in AMPA Receptor-Auxiliary Subunit Complexes. Neuron 94, 569–580.e5 (2017).

37. Hale, W. D. et al. Allosteric competition and inhibition in AMPA receptors. Nat. Struct. Mol. Biol. 1–11 (2024) doi:10.1038/s41594-024-01328-0.

38. Hale, W. D. et al. Structure of transmembrane AMPA receptor regulatory protein subunit γ2. Nat. Commun. 16, 671 (2025).

39. Yelshanskaya, M. V., Patel, D. S., Kottke, C. M., Kurnikova, M. G. & Sobolevsky, A. I. Opening of glutamate receptor channel to subconductance levels. Nature 605, 172–178 (2022).

40. Zhang, D. et al. Structural mobility tunes signalling of the GluA1 AMPA glutamate receptor. Nature 621, 877–882 (2023).

41. Rossmann, M. et al. Subunit-selective N-terminal domain associations organize the formation of AMPA receptor heteromers. EMBO J. 30, 959–971 (2011).

42. Dutta, A., Shrivastava, I. H., Sukumaran, M., Greger, I. H. & Bahar, I. Comparative Dynamics of NMDA- and AMPA-Glutamate Receptor N-Terminal Domains. Structure 20, 1838–1849 (2012).

43. Kumar Mondal, A., Carrillo, E., Jayaraman, V. & Twomey, E. C. Glutamate gating of AMPA-subtype iGluRs at physiological temperatures. Nature 641, 788–796 (2025).

44. Smart, O. S., Neduvelil, J. G., Wang, X., Wallace, B. A. & Sansom, M. S. P. HOLE: A program for the analysis of the pore dimensions of ion channel structural models. J. Mol. Graph. 14, 354–360 (1996).

45. Rossmann, M. et al. Subunit-selective N-terminal domain associations organize the formation of AMPA receptor heteromers. EMBO J. 30, 959–971 (2011).

46. Smith, T. C., Wang, L.-Y. & Howe, J. R. Heterogeneous Conductance Levels of Native AMPA Receptors. J. Neurosci. 20, 2073–2085 (2000).

47. Prieto, M. L. & Wollmuth, L. P. Gating modes in AMPA receptors. J. Neurosci. Off. J. Soc. Neurosci. 30, 4449–4459 (2010).

48. Smith, T. C. & Howe, J. R. Concentration-dependent substate behavior of native AMPA receptors. Nat. Neurosci. 3, 992–997 (2000).

49. Rosenmund, C., Stern-Bach, Y. & Stevens, C. F. The tetrameric structure of a glutamate receptor channel. Science 280, 1596–1599 (1998).

50. Poon, K., Nowak, L. M. & Oswald, R. E. Characterizing Single-Channel Behavior of GluA3 Receptors. Biophys. J. 99, 1437–1446 (2010).

51. Shi, E. Y. et al. Noncompetitive antagonists induce cooperative AMPA receptor channel gating. J. Gen. Physiol. 151, 156–173 (2019).

52. Yuan, C. L. et al. Modulation of AMPA Receptor Gating by the Anticonvulsant Drug, Perampanel. ACS Med. Chem. Lett. 10, 237–242 (2019).

53. Robert, A. & Howe, J. R. How AMPA Receptor Desensitization Depends on Receptor Occupancy. J. Neurosci. 23, 847–858 (2003).

54. Zhang, W., Cho, Y., Lolis, E. & Howe, J. R. Structural and Single-Channel Results Indicate That the Rates of Ligand Binding Domain Closing and Opening Directly Impact AMPA Receptor Gating. J. Neurosci. 28, 932–943 (2008).

55. Punjani, A., Rubinstein, J. L., Fleet, D. J. & Brubaker, M. A. cryoSPARC: algorithms for rapid unsupervised cryo-EM structure determination. Nat. Methods 14, 290–296 (2017).

56. Pettersen, E. F. et al. UCSF ChimeraX: Structure visualization for researchers, educators, and developers. Protein Sci. 30, 70–82 (2021).

57. Emsley, P. & Cowtan, K. Coot: model-building tools for molecular graphics. Acta Crystallogr. D Biol. Crystallogr. 60, 2126–2132 (2004).

58. Croll, T. I. ISOLDE: a physically realistic environment for model building into low-resolution electron-density maps. Acta Crystallogr. Sect. Struct. Biol. 74, 519–530 (2018).

59. Liebschner, D. et al. Macromolecular structure determination using X-rays, neutrons and electrons: recent developments in Phenix. Acta Crystallogr. Sect. Struct. Biol. 75, 861–877 (2019).

60. Morin, A. et al. Collaboration gets the most out of software. eLife 2, e01456 (2013).

61. Williams, C. J. et al. MolProbity: More and better reference data for improved all-atom structure validation. Protein Sci. 27, 293–315 (2018).

