## Extended Data Table 1 for "Architecture, Activation, and Conformational Plasticity in the GluA4 AMPA Receptor"

### Cryo-EM data collection, refinement and validation statistics

| | GluA4/TARPγ2 (EMDB-xxxx) (PDB xxxx) | Substate 1 (EMDB-xxxx) (PDB xxxx) | Substate 2 (EMDB-xxxx) (PDB xxxx) | Substate 3 (EMDB-xxxx) (PDB xxxx) | Substate 4 (EMDB-xxxx) (PDB xxxx) | Substate 5 (EMDB-xxxx) (PDB xxxx) |
|---|---|---|---|---|---|---|
| **Data collection and processing** | | | | | | |
| Magnification | 130,000x | 130,000x | 130,000x | 130,000x | 130,000x | 130,000x |
| Voltage (kV) | 300 | 300 | 300 | 300 | 300 | 300 |
| Electron exposure (e–/Å$^2$) | 40 | 40 | 40 | 40 | 40 | 40 |
| Defocus range (μm) | -1.0-2.5 | -1.0-2.5 | -1.0-2.5 | -1.0-2.5 | -1.0-2.5 | -1.0-2.5 |
| Pixel size (Å) | 0.93 | 0.93 | 0.93 | 0.93 | 0.93 | 0.93 |
| Symmetry imposed | N/A | N/A | N/A | N/A | N/A | N/A |
| Initial particle images (no.) | 2,418,306 | 2,418,306 | 2,418,306 | 2,418,306 | 2,418,306 | 2,418,306 |
| Final particle images (no.) | 253,513 | 21,063 | 56,205 | 82,102 | 59,237 | 22,575 |
| Map resolution (Å) FSC = 0.143 | 3.31 | 4.01 | 3.82 | 3.82 | 3.89 | 4.06 |
| Map resolution range (Å) | 2-42 | 2.4-50 | 2.2-47 | 2.5-45 | 2.6-49 | 2.2-50 |
| **Refinement** | | | | | | |
| Initial model used (PDB code) | 5WEO | GluA4/TARPγ2 | GluA4/TARPγ2 | GluA4/TARPγ2 | GluA4/TARPγ2 | GluA4/TARPγ2 |
| Model resolution (Å) FSC threshold = 0.143 | 3.27 | 4.2 | 4.1 | 4.1 | 4.1 | 4.2 |
| Map sharpening *B* factor (Å$^2$) | 92.6 | 49.0 | 67.4 | 82.4 | 71.6 | 56.2 |
| Model composition | | | | | | |
| Non-hydrogen atoms | 18,108 | 18,108 | 18,108 | 18,108 | 18,108 | 18,108 |
| Protein residues | 2308 | 2308 | 2308 | 2308 | 2308 | 2308 |
| Ligands | 4 | 4 | 4 | 4 | 4 | 4 |
| *B* factors (Å$^2$) | | | | | | |
| Protein | 23.68/201.32/104.60 | 73.31/241.79/144.22 | 35.46/219.60/116.58 | 31.74/182.41/108.13 | 30.03/187.09/96.45 | 41.06/253.90/126.65 |
| Ligand | 78.83/135.24/101.57 | 164.77/178.55/169.39 | 126.66/131.31/128.81 | 128.98/140.55/134.79 | 98.64/104.73/102.51 | 136.72/143.45/140.18 |
| R.m.s. deviations | | | | | | |
| Bond lengths (Å) | 0.003 | 0.003 | 0.004 | 0.003 | 0.002 | 0.002 |
| Bond angles (°) | 0.536 | 0.502 | 0.526 | 0.481 | 0.460 | 0.759 |
| **Validation** | | | | | | |
| MolProbity score | 1.53 | 1.7 | 1.73 | 1.65 | 1.57 | 1.55 |
| Clashscore | 4.35 | 5.39 | 6.00 | 5.20 | 4.54 | 4.98 |
| Poor rotamers (%) | 0.00 | 0.00 | 0.00 | 0.00 | 0.00 | 0.00 |
| Ramachandran plot | | | | | | |
| Favored (%) | 95.43 | 93.84 | 94.02 | 94.64 | 95.08 | 95.79 |
| Allowed (%) | 4.43 | 6.03 | 5.90 | 5.10 | 4.74 | 3.99 |
| Disallowed (%) | 0.13 | 0.13 | 0.09 | 0.27 | 0.18 | 0.22 |
